# The profiles of mammals, birds and fish in twelve water bodies on the Island of Montreal by mitochondrial DNA: detection of invasive animals and meat and fish waste

**DOI:** 10.1101/2025.05.16.654469

**Authors:** Richard Villemur

**Affiliations:** INRS-Centre Armand-Frappier Santé Biotechnologie, Laval (Québec) H7V 1B7

## Abstract

The Island of Montreal is composed of a variety of environments, such as undeveloped natural areas, urban parks, and dense residential areas. These environments are home to several types of surface waters, such as ponds, marshes, lakes, streams, and rivers, where some wild animals have adapted. Among them are exotic and invasive animals, which could displace natural animals or endanger people through unwanted interactions. The Montreal Urban Community (MUC) wishes to assess the extent of the presence of exotic or invasive animals in waterbodies on the Island of Montreal to coordinate mitigation measures. Here, I present animal profiles, primarily mammals, birds, and fish, from twelve waterbodies on the Island of Montreal using amplicon sequencing of mitochondrial DNA from environmental DNA. Water samples were collected, and total DNA was extracted. PCR amplifications of a portion of the mitochondrial 16S ribosomal RNA genome were performed with primers designed to primarily target mammals, fish, and birds. After sequencing, the animal lineages and their proportions were determined. Our results showed that invasive animals, primarily fish such as pumpkinseed, common carp or brown bullhead, were detected in high proportions in five water bodies. However, I did not detect any exotic animals. I also detect significant proportions of sequences affiliated with bovine, porcine and several marine fish species, which was unexpected as these animals are not living on the Island of Montreal. Given the commercial or industrial activities surrounding the water bodies, sequences of these unexpected animals are likely related to the release of meat or seafood waste into the water bodies. Our molecular approach is effective for assessing the animal profiles of water bodies. However, caution should be exercised when interpreting the presence of animal waste.

## Introduction

Urbanized areas include a variety of environments, such as undeveloped natural areas, city parks, and dense residential areas. These environments support several types of surface waters, such as ponds, marshes, lakes, streams, and rivers. Besides humans, domestic animals, and livestock if farms are nearby, some wild animals have adapted to urbanized areas. Additionally, there are internal city reports indicating the exotic animals acquired by citizens are intentionally release into the water bodies or escape from their owners; some survive, while others cannot survive the winter. For example, exotic aquarium fish or amphibians (e.g., turtles) may be present in water bodies. Additionally, the migration of predatory animals (e.g., coyotes) into urban areas has also been observed. The presence of exotic or invasive animals can endanger natural wildlife by displacing naturally occurring animals or endanger people through unwanted interactions. Faced with these potential problems, the Montreal Urban Community (CUM) wanted to assess the extent of the presence of exotic or invasive animals in water bodies on the Island of Montreal in order to coordinate mitigation measures.

Mitochondrial DNA (mtDNA) has become the standard barcoding tool for identifying the presence of animal species in different environments. We have developed PCR primers to target the mtDNA of the majority of mammals, birds, and fish, and to some extent reptiles and amphibians (Ragot & Villemur, 2022). These primers amplify a region of the mitochondrial genome (gene encoding part of the mitochondrial 16S rRNA gene) approximately 250–300 nt in length (amplicon). By using these primers to PCR amplify mtDNA from environmental DNA (eDNA), we generate an amplicon composed of different sequences representative of mammals, birds, and fish. This amplicon is then sequenced using next-generation sequencing technologies (e.g., Illumina). In a previous study, we used these primers to detect the presence of these animals and determine their proportions in two watersheds (86 sampling sites) (Ragot *et al.*, 2023), and in the outlet of a stream located in an agricultural area during the summer (Villemur, 2025) in order to assess the potential contribution of fecal contamination of surface water by multiple animals.

Using the same approach, we monitored twelve water bodies (ponds, marshes, streams) at different locations on the Island of Montreal in September 2022 to detect the presence of mtDNA sequences primarily associated with mammals, fish, and birds. The objective of this monitoring is to observe the presence of exotic and invasive animals and estimate their proportions. Our results showed that invasive animals, primarily fish, were detected, but not exotic animals. Our approach revealed interesting observations: the detection in relatively high levels of sequences affiliated with bovine, or porcine, or to animals unlikely to live in the region, such as marine fish, suggesting the release of meat and fish waste into the water bodies.

## Material and Methods

### Water collection

Twelve sampling sites were identified by the MUC (Fig. 1, Table 1). MUC representatives, Jean-Philippe Lafleur and Suzanne Boulet (*Réseau de suivi du milieu aquatique; Service de l’environnement; Division du contrôle des rejets et suivi environnemental*), facilitated access to the sites. Most of these sites were located in parks or streams surrounding dense urban areas. Water samples were collected on September 18 (samples 1 to 6) and 25 (samples 7 to 12), 2022. Water samples (1 L) were collected and placed on ice. They were filtered through 0.45 µM filters on the same day. The filters were stored at -70 °C until use. On September 18, heavy rain occurred during sample collection at some sites. The temperature was 11°C on September 18 and 15°C on September 25.

**Table 1.** Simpling location sites, with corresponding number of sequenced reads.

| Sample name | Location | Latitude (N) | Longitude (W) | Number of reads |
| --- | --- | --- | --- | --- |
| Mtl01 | Lake *Centenaire* William Cosgrove Park | 45.488608 | 73.810611 | 238877 |
| Mtl02 | Marsh *parc-nature du Boisé-de-l'Île-Bizard* | 45.513459 | 73.889475 | 271748 |
| Mtl03 | Stream *O'Connell* | 45.460540 | 73.907099 | 367450 |
| Mtl04 | River *à l'Orme* | 45.444474 | 73.915883 | 195994 |
| Mtl05 | Stream *Denis* (manhole) | 45.448383 | 73.779922 | 255770 |
| Mtl06 | Stream *Bouchard* | 45.440807 | 73.726426 | 275267 |
| Mtl07 | Basin *de La Brunante* | 45.509845 | 73.706756 | 266814 |
| Mtl08 | Stream *Bertrand* | 45.508726 | 73.758132 | 359966 |
| Mtl09 | Pond *Angrignon* Park | 45.444301 | 73.596931 | 327660 |
| Mtl10 | Lake *Lacoursière* Park | 45.465688 | 73.549828 | 335560 |
| Mtl11 | Stream *De Montigny* | 45.613607 | 73.599982 | 344046 |
| Mtl12 | Marsh *parc-nature de la Pointe-aux-Prairies* | 45.692166 | 73.522569 | 358498 |

Reads were derived from amplicon sequencing of PCR-amplified mtDNA of eDNA extracted from samples.

**Figure 1.**
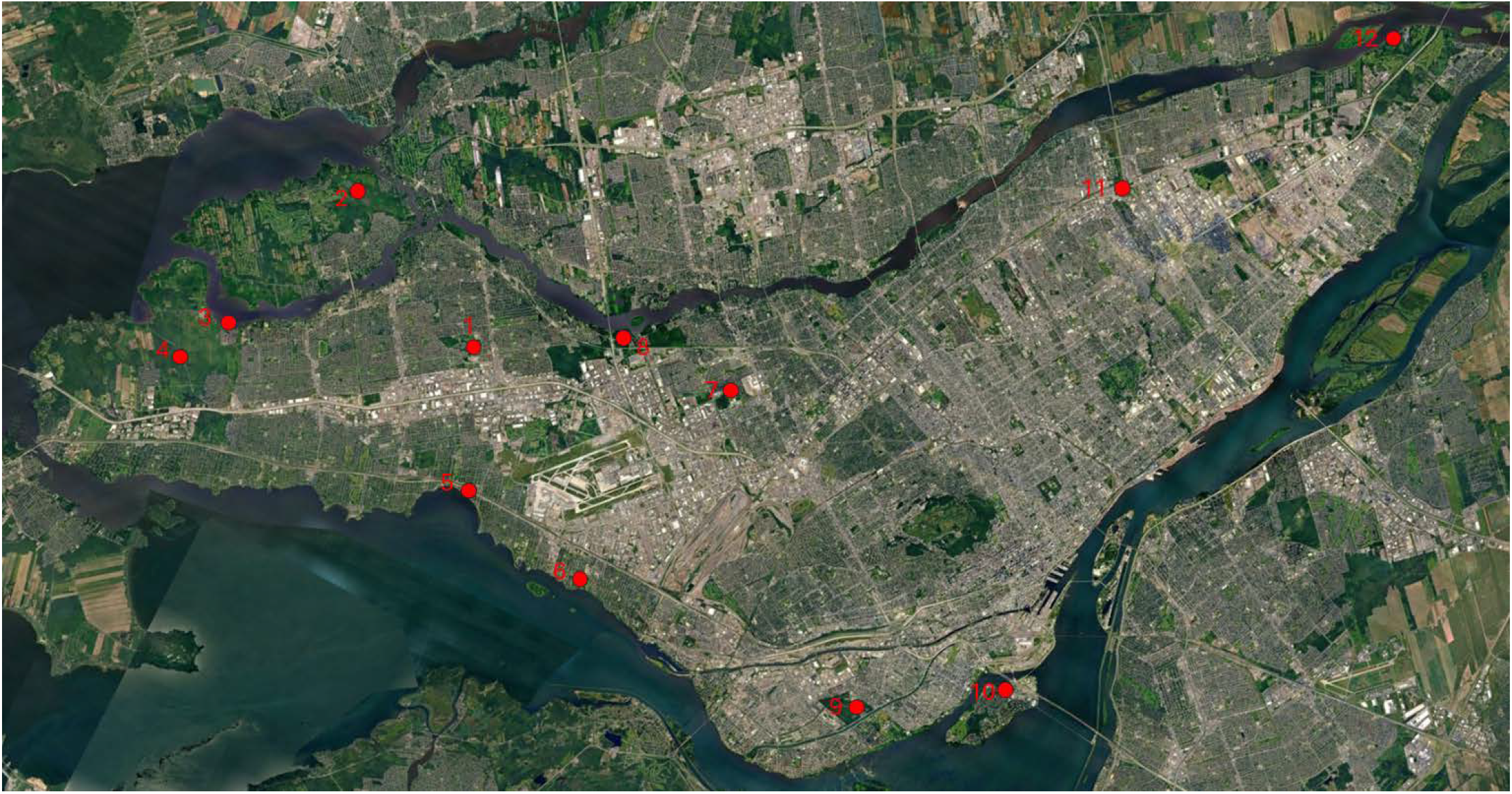
Island of Montreal with the location of the sampling sites (in red). See Table 1 for details.

### DNA extraction and libraries

DNA extraction from filters, and the generation of libraries for amplicon sequencing were performed at the Villemur’s laboratory as described (Ragot & Villemur, 2022), with modifications described in Villemur (2025). Raw sequencing data were deposited in Sequence Read Archive (SRA) at the National Center for Biotechnology Information (NCBI: https://www.ncbi.nlm.nih.gov/),Bioproject ID PRJNA1263451, Biosample accessions SAMN48521554 to SAMN48521565.

The identified organisms were listed by their Latin scientific names. The correspondence of these names to the animal names is available in the Appendix, after the References section.

## Results

### Site Mtl01: Lake of the *Centenaire William Cosgrove* Park (Table 2)

This is a small lake located in a park in a dense urban area. The sample was collected at the mouth of the lake, at low flow. Fish mtDNA sequences accounted for 75% of the sequences. The proportions of squirrel (*Sciurus carolinensis*) and human sequences were 13% and 10%, respectively. Among fish, the most abundant sequences were those of pumpkinseed (*Lepomis gibbosus*; 30%) and common carp (*Cyprinus carpio*; 10%). These fish are usually found in ponds but could also have been introduced by people. They are considered invasive species. Poultry (*Gallus gallus*) and porcine (*Sus scrofa*) mtDNA sequences were fewer in proportion than those of humans (see discussion).

**Table 2.** Affiliated organisms found in the Mtl01, Mtl02 and Mtl03 sampling sites by mtDNA sequencing.

| Mtl01 | % | Mtl02 | % | Mtl03 | % |
| --- | --- | --- | --- | --- | --- |
| *Homo sapiens* | 9.94 | *Homo sapiens* | 5.90 | *Homo sapiens* | 1.09 |
| *Gallus gallus* | 0.451 | *Sus scrofa* | 0.0824 | *Ovis aries* | 0.111 |
| *Sus scrofa* | 0.187 | *Bos taurus* | 0.0473 | *Sus scrofa* | 0.0638 |
| *Sciurus carolinensis* | 13.21 | *Ondatra zibethicus* | 0.192 | *Bos taurus* | 0.0024 |
| *Tamias striatus* | 0.211 | *Lithobates catesbeianus* | 0.0922 | *Sylvilagus floridanus* | 1.60 |
| *Ondatra zibethicus* | 0.0036 | *Branta canadensis* | 8.49 | *Ondatra zibethicus* | 0.213 |
| *Sturnus vulgaris* | 0.536 | *Anas platyrhynchos* | 1.35 | *Procyon lotor* | 0.162 |
| *Passer montanus* | 0.107 | *Ameiurus nebulosus* | 26.25 | *Peromyscus leucopus* | 0.0173 |
| *Branta canadensis* | 0.0414 | *Pimephales promelas* | 24.03 | *Sciurus carolinensis* | 0.0036 |
| *Pimephales promelas* | 31.98 | *Lepomis gibbosus* | 18.39 | *Chelydra serpentina* | 0.0045 |
| *Lepomis gibbosus* | 30.40 | *Phoxinus eos* | 7.55 | *Anas platyrhynchos* | 0.0013 |
| *Cyprinus carpio* | 9.83 | *Culaea inconstans* | 3.56 | *Ameiurus nebulosus* | 21.96 |
| *Ameiurus nebulosus* | 1.36 | *Notemigonus crysoleucas* | 2.39 | *Culaea inconstans* | 2.70 |
| *Perca flavescens* | 0.564 | *Hybognathus spp.* | 0.365 | *Notemigonus crysoleucas* | 1.24 |
| *Ambloplites rupestris* | 0.425 | *Phoxinus neogaeus* | 0.149 | *Catostomus spp.* | 1.19 |
| *Notemigonus crysoleucas* | 0.375 | *Catostomus spp.* | 0.0129 | *Pimephales promelas* | 0.862 |
| *Culaea inconstans* | 0.0589 | *Cyprinus carpio* | 0.0016 | *Cyprinella spiloptera* | 0.853 |
| *Catostomus spp.* | 0.0089 | *Umbra limi* | 0.0016 | *Lepomis gibbosus* | 0.846 |
| *Esox lucius* | 0.0066 | *Plumatella* | 0.0554 | *Fundulus diaphanus* | 0.741 |
| *Bosmina liederi* | 0.158 | *Tetragoneuria cynosura* | 0.0088 | *Tinca tinca* | 0.650 |
| *Bosmina freyi* | 0.102 | *Simocephalus serrulatus* | 0.0036 | *Pimephales notatus* | 0.249 |
| *Daphnia parvula* | 0.0174 | *Dioscorea rotundata* | 0.0004 | *Carassius auratus* | 0.0576 |
| *Plumatella emarginata* | 0.0153 |  |  | *Etheostoma olmstedii* | 0.0611 |
|  |  |  |  | *Ambloplites rupestris* | 0.0595 |
|  |  |  |  | *Semotilus atromaculatus* | 0.0022 |
|  |  |  |  | *Micropterus spp.* | 0.0014 |
|  |  |  |  | *Bosmina freyi* | 65.16 |
|  |  |  |  | *Plumatella emarginata* | 0.105 |

...: human, pet and livestock/meat. ...: Wild mammals. ...: birds. ...: amphibians, reptile. ...: fish.
...: others such as cretaceans, insects.

### Site Mtl02: Marsh *parc-nature du Boisé-de-l’Île-Bizard* (Table 2)

This is a park located in a semi-urban area surrounded by forest with houses relatively far from the marsh (1 km). The lake was relatively clear with many aquatic plants. It was raining (a light rain). The sample was collected in the middle of the lake; the water was stagnant at the sampling site. As at site Mtl01, the proportion of fish sequences was 75%. Sequences affiliated with large fish such as the brown bullhead (*Ameiurus nebulosus*) and pumpkinseed were present in high proportions (26.3% and 18.4%, respectively). We observed several bird species. Sequences affiliated with the Canada goose (*Branta canadensis*) represented 8.5% of the total sequences. The proportion of human sequences was close to 6%. Sequences related to porcine and bovine (*Bos taurus*), as well as wild animals such as the muskrat (*Ondatra zibethicus*) were present in low proportion (<1%).

### Site Mtl03 : Stream *O’Connell* (Table 2)

This stream is the outlet of a small pond located in a dense urban area. At the time of sampling, the stream was experiencing high flow, likely related to heavy rainfall during the day. The sampling site was located near a boulevard. The most relative abundant mtDNA sequences (65.2%) were related to a crustacean, possibly affiliated with the genus *Bosmina*. For example, *Bosmina freyi* is an arthropod species in the family Bosminidae. These species are associated with freshwater habitats. They are herbivores (https://eol.org/pages/1039335). Their presence could be related to the degradation of organic matter due to water stagnation in the pond.

Sequences related to the brown bullhead, an invasive fish species, were found in 22% of cases. Human sequences (1.1%), as well as sequences from ovine (*Ovis aries*), porcine and bovine, were found. Sequences affiliated with the wild mammals rabbit (*Sylvilagus floridanus*), muskrat, and white-footed mouse (*Peromyscus leucopus*) were observed in low proportion.

### Mtl04 : River *à l’Orme* (Table 3)

The sampling site was located in a wooded environment, few kilometers from urban and commercial areas where the river passes upstream. During sampling, pouring rain occurred, and the water flow was very high. mtDNA sequences affiliated with the small-mouth stickleback (*Culaea inconstans*) were very high in proportion (69.1%). We observed mtDNA sequences affiliated with many wild animals normally found in forests South of the Province of Quebec. We also noted the presence of sequences affiliated with the black bear (*Ursus americanus*; 0.063%). A captive bear was present nearby in a small zoo. Human-affiliated sequences were the second highest in proportion (6.1%). Sequences affiliated with bovine, cat (*Felis silvestris*), porcine, dog (*Canis lupus*), poultry and Atlantic salmon (*Salmo salar*; see Discussion) were found in low proportion.

**Table 3.** Affiliated organisms found in the Mtl04, Mtl05 and Mtl06 sampling sites by mtDNA sequencing.

| Mtl04 | % | Mtl05 | % | Mtl06 | % |
| --- | --- | --- | --- | --- | --- |
| *Homo sapiens* | 6.07 | *Homo sapiens* | 2.38 | *Homo sapiens* | 2.94 |
| *Bos taurus* | 0.245 | *Canis lupus* | 0.209 | *Gallus gallus* | 0.363 |
| *Felis silvestris* | 0.193 | *Bos taurus* | 0.194 | *Bos taurus* | 0.282 |
| *Sus scrofa* | 0.122 | *Gallus gallus* | 0.0123 | *Canis lupus* | 0.178 |
| *Canis lupus* | 0.0661 | *Sus scrofa* | 0.0094 | *Sus scrofa* | 0.155 |
| *Gallus gallus* | 0.0319 | *Procyon lotor* | 59.99 | *Felis catus* | 0.132 |
| *Salmo salar* | 0.0102 | *Sylvilagus floridanus* | 23.70 | *Sciurus carolinensis* | 1.05 |
| *Sciurus carolinensis* | 4.49 | *Sciurus carolinensis* | 2.85 | *Ondatra zibethicus* | 0.908 |
| *Microtus spp.* | 3.98 | *Peromyscus leucopus* | 0.990 | *Procyon lotor* | 0.891 |
| *Mephitis mephitis* | 1.96 | *Microtus spp.* | 0.159 | *Microtus spp.* | 0.559 |
| *Sylvilagus floridanus* | 0.830 | *Mephitis mephitis* | 0.110 | *Rattus norvegicus* | 0.223 |
| *Procyon lotor* | 0.677 | *Rattus norvegicus* | 0.0612 | *Mephitis mephitis* | 0.207 |
| *Peromyscus leucopus* | 0.467 | *Ondatra zibethicus* | 0.0893 | *Sylvilagus floridanus* | 0.196 |
| *Blarina brevicauda* | 0.409 | *Blarina brevicauda* | 0.0410 | *Peromyscus leucopus* | 0.195 |
| *Ondatra zibethicus* | 0.407 | *Tamias striatus* | 0.0422 | *Sciurus niger* | 0.112 |
| *Vulpes vulpes* | 0.255 | *Sciurus niger* | 0.0260 | *Marmota monax* | 0.0817 |
| *Tamias striatus* | 0.166 | *Mus musculus* | 0.0244 | *Mus musculus* | 0.0352 |
| *Marmota monax* | 0.103 | *Odocoileus virginianus* | 0.0197 | *Columba livia* | 48.92 |
| *Rattus norvegicus* | 0.101 | *Marmota monax* | 0.0182 | *Anas crecca* | 1.26 |
| *Condylura cristata* | 0.0865 | *Tamiasciurus hudsonicus* | 0.0064 | *Turdus spp.* | 1.02 |
| *Tamiasciurus hudsonicus* | 0.0633 | *Alces alces* | 0.0029 | *Larus spp.* | 0.644 |
| *Ursus americanus* | 0.0628 | *Vulpes vulpes* | 0.0035 | *Sturnus vulgaris* | 0.317 |
| *Didelphis virginiana* | 0.0342 | *Columba livia* | 0.311 | *Melospiza melodia* | 0.177 |
| *Meleagris gallopavo* | 0.0240 | *Sturnus spp.* | 0.160 | *Spizella passerina* | 0.145 |
| *Lepus spp.* | 0.0235 | *Corvus brachyrhynchos* | 0.0672 | *Vermivora celata* | 0.120 |
| *Odocoileus virginianus* | 0.0023 | *Larus spp.* | 0.0544 | *Passer montanus* | 0.0710 |
| *Anas platyrhynchos* | 0.328 | *Anas platyrhynchos* | 0.0430 | *Anaxyrus spp.* | 0.426 |
| *Turdus spp.* | 0.165 | *Turdus spp.* | 0.0420 | *Pseudacris crucifer* | 0.0798 |
| *Aphelocoma* | 0.0607 | *Quiscalus quiscula* | 0.0266 | *Lithobates pipiens* | 0.0238 |
| *Larus spp.* | 0.0372 | *Branta canadensis* | 0.0223 | *Hyla spp.* | 0.0156 |
| *Aix galericulata* | 0.0304 | *Dendroica coronata* | 0.0139 | *Storeria dekayi* | 0.0114 |
| *Carduelis tristis* | 0.0298 | *Spizella passerina* | 0.0127 | *Culaea inconstans* | 2.27 |
| *Melospiza melodia* | 0.0219 | *Zenaida macroura* | 0.0064 | *Semotilus atromaculatus* | 1.66 |
| *Charadrius vociferus* | 0.0189 | *Meleagris gallopavo* | 0.0025 | *Catostomus spp.* | 1.36 |
| *Regulus calendula* | 0.0099 | *Passer montanus* | 0.0037 | *Ictiobus bubalus* | 0.470 |
| *Catharus fuscescens* | 0.0038 | *Falco sparverius* | 0.0014 | *Pimephales promelas* | 0.245 |
| *Columba livia* | 0.0069 | *Picidae* | 0.0051 | *Luxilus spp.* | 0.134 |
| *Zonotrichia spp.* | 0.0043 | *Anaxyrus spp.* | 7.56 | *Mugil cephalus* | 0.131 |
| *Dumetella carolinensis* | 0.0026 | *Pseudacris crucifer* | 0.0311 | *Carassius auratus* | 0.0193 |
| *Anaxyrus spp.* | 3.90 | *Ambystoma unisexual* | 0.0043 | *Rhomboplites aurorubens* | 12.07 |
| *Pseudacris crucifer* | 1.45 | *Lithobates catesbeianus* | 0.0012 | *Salmo salar* | 7.11 |
| *Notophthalmus spp.* | 0.117 | *Pimephales promelas* | 0.190 | *Scomberomorus cavalla* | 2.99 |
| *Plethodon cinereus* | 0.111 | *Semotilus atromaculatus* | 0.181 | *Pomatomus saltatrix* | 1.63 |
| *Rana pipiens* | 0.0472 | *Umbra limi* | 0.0158 | *Dicentrarchus labrax* | 1.35 |
| *Lithobates clamitans* | 0.0186 | *Cyprinella spiloptera* | 0.109 | *Scomber spp.* | 1.25 |
| *Culaea inconstans* | 69.13 | *Culaea inconstans* | 0.096 | *Calamus penna* | 1.03 |
| *Cyprinella spiloptera* | 2.46 | *Catostomus spp.* | 0.0893 | *Sarotherodon galilaeus* | 1.01 |
| *Esox lucius* | 0.494 | *Lepomis gibbosus* | 0.0029 | *Leucoraja erinacea* | 0.535 |
| *Umbra limi* | 0.247 | *Bosmina freyi* | 0.0087 | *Caranx hippos et autres* | 0.389 |
| *Ameiurus nebulosus* | 0.169 | *Forficula auricularia* | 0.0023 | *Euthynnus alletteratus* | 0.345 |
| *Cottus bairdii* | 0.0615 |  |  | *Merluccius spp.* | 0.308 |
| *Semotilus atromaculatus* | 0.0410 |  |  | *Seriola* | 0.282 |
| *Phoxinus neogaeus* | 0.0299 |  |  | *Epinephelus flavolimbatus* | 0.269 |
| *Catostomus spp.* | 0.0114 |  |  | *Carcharhinus* | 0.226 |
| *Pimephales notatus* | 0.0163 |  |  | *Xiphias gladius* | 0.223 |
| *Bosmina freyi* | 0.0742 |  |  | *Pagrus pagrus* | 0.218 |
| *Plumatella repens* | 0.0281 |  |  | *Scomberomorus spp.* | 0.196 |
|  |  |  |  | *Sparus aurata* | 0.195 |
|  |  |  |  | *Paralichthys dentatus* | 0.155 |
|  |  |  |  | *Morone chrysops* | 0.106 |
|  |  |  |  | *Sander vitreus* | 0.0385 |
|  |  |  |  | *Aporrectodea rosea* | 0.0430 |
|  |  |  |  | *Plumatella* | 0.0276 |

...: human, pet and livestock/meat. ...: Wild mammals. ...: birds. ...: amphibians, reptile. ...: fish. ...: others such as cretaceans, insects. ...: Mainly marine fish suspected to originate from fish wholesalers nearby.

### Mtl05 : Stream *Denis* (Table 3)

Sampling was conducted in a manhole where the stream passes under the street. According to the Google Earth map, the stream originates in an industrial area, flows through a vacant lot surrounded by a golf course and housing, before reaching a pipe that passes under a highway, then under urban streets, to the St. Lawrence River. High proportions of sequences affiliated with raccoon (60%; *Procyon lotor*) and rabbit (23.7%) were found, suggesting the presence of many members of these species near the stream. Human-affiliated sequences (2.4%) and low proportions of sequences affiliated with dog, bovine, poultry, porcine, and moose (*Alces alces*; see discussion) were found.

### Mtl06 : Stream *Bouchard* (Table 3)

The sample was collected under a bridge crossing the stream, just before its mouth into the St. Lawrence River. We did not expect to find 31.8% of sequences associated with seafood. From downstream to upstream, the stream crosses a residential area, then a commercial area, passes under a highway, and then crosses an industrial area, where we notably noticed fish and seafood wholesalers where the stream passes right behind. The presence of these wholesalers could explain the great diversity of seafood-related species, such as shark (*Carcharhinus*), swordfish (*Xiphias gladius*) and mackerel (*Scomberomorus*). Their website, where seafood is sold, specifically reveals the fish detected in the stream. It is possible that the presence of these fish is due to faulty sewer pipes that have discharged fish waste directly into the stream.

The high proportion of pigeons (*Columba livia*; 48.9%) can be attributed to water sampling at the bridge, where they often roost. Another possibility is that they are attracted to fish waste discharged into the stream. Human-related sequences (3.9%) and a small proportion of sequences affiliated with bovine, dog, porcine, and cat were found.

### Mtl07 : Basin *de La Brunante* (Table 4)

This small basin is located in a dense urban area. The water was stagnant, and fish and birds were visible. Sequences affiliated with the invasive fish pumpkinseed (42.9%), common carp (8.5%), brown bullhead (1.4%), and goldfish (*Carassius auratus*; 3.5%) were found. A high proportion of sequences (35.9%) affiliated with the crustaceans *Ceriodaphnia* or *Daphnia* were also found. Daphnia are naturally present in stagnant waters and associated with the degradation of organic matter in aquatic bodies such as leaf litter. Humans accounted for 2.9% of the sequences, with a low proportion of sequences affiliated with bovine (0.6%).

**Table 4.**
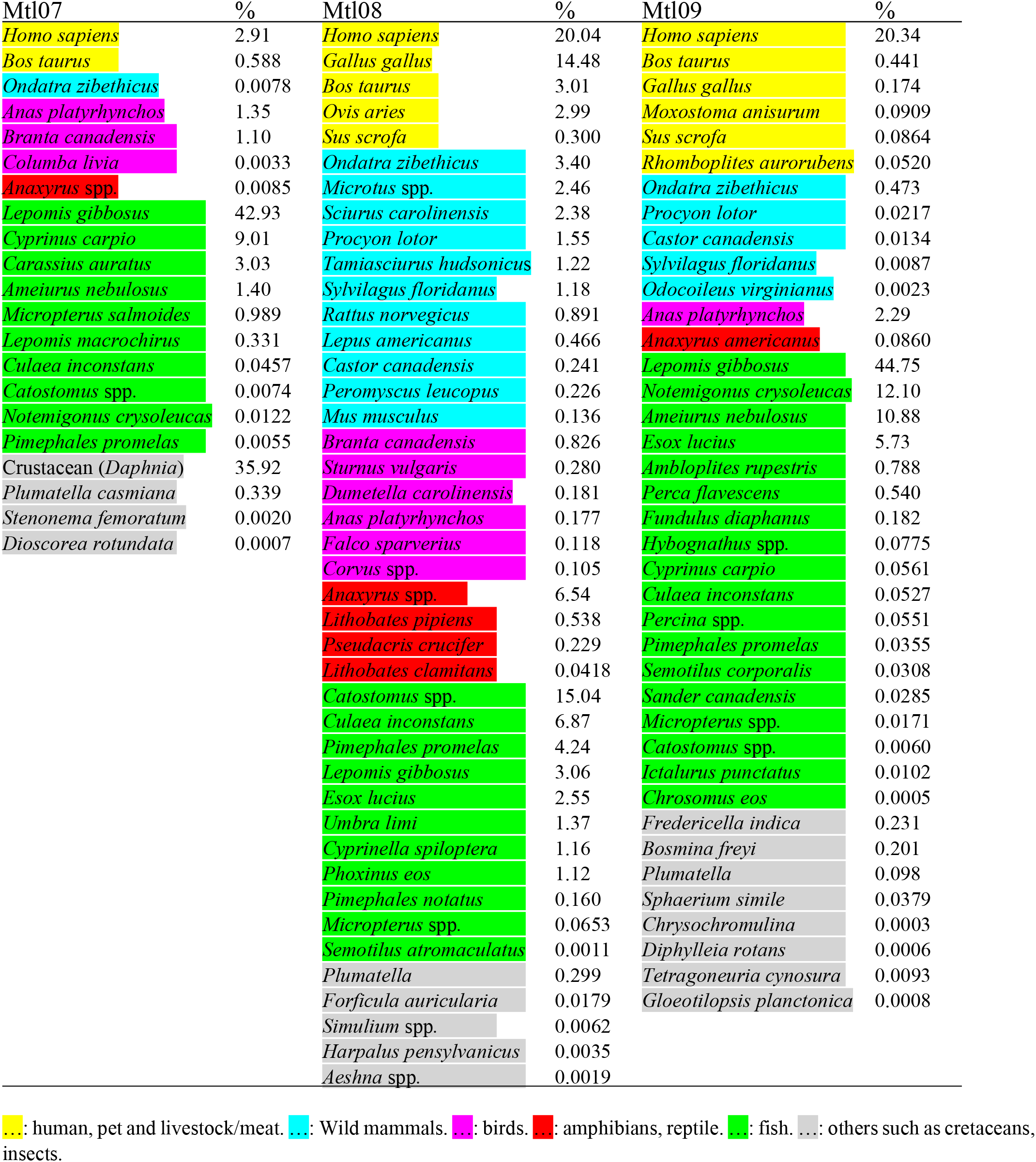
Affiliated organisms found in the Mtl07, Mtl08 and Mtl09 sampling sites by mtDNA sequencing.

### Mtl08: Stream *Bertrand* (Table 4)

The stream flows through a large, mostly wooded municipal park and then runs alongside a residential area. Upstream, the stream is very shallow and flows through an industrial area. Water flow was low. The sampling site was located along a boulevard near an urban area, which could explain the high proportion of human-related mtDNA sequences (20%). Faulty wastewater pipe connections could explain the high proportion of human sequences. There was also a high proportion of sequences affiliated with poultry (14.5%), bovine (3%), and ovine (3%), which may not be exclusively explained by the presence of undigested meat in human feces. Two takeout restaurants are located upstream near the stream. As with the River *à l’Orme* site, sequences related to several wild mammals, amphibians, and birds normally found in the southern part of the Province of Quebec were found.

### Mtl09 : Pond of the *Angrignon* Park (Table 4)

This pond is located in a park, surrounded by a dense urban area. Citizens reported observing turtles (possibly exotic animals) a little further west of the pond. However, sampling there was not possible, and we had to collect samples where the water was stagnant. Unfortunately, no turtle-related sequences were found, highlighting the difficulty of choosing the right sampling location. It would probably have been preferable to sample at the outlet of the pond, where water mixing occurred. A high proportion of sequences related to the invasive fish pumpkinseed (44.8%), brown bullhead (12.1%), and northern pike (*Esox lucius*; 5.7%) were found. Human sequences were second in proportion (20.3%), followed by a small proportion of sequences from bovine, poultry, ovine and vermillion snapper (*Rhomboplites aurorubens*; see discussion).

### Mtl10 : Lake of Park *Lacoursière* (Table 5)

This is a small pond with stagnant water. It is located in a dense urban area. Despite this environment, the site had the lowest proportion of human-affiliated mtDNA sequences (0.8%) and very low proportions of sequences affiliated with bovine, poultry or porcine. It is possible that this new urban area has better wastewater connection infrastructure. The stagnant water appears to have favored the small crustacean *Cypridopsis vidua* (29%), which is frequently found in these type of water bodies. A high proportion of sequences affiliated with the invasive fish brown bullhead was found (13.7%).

**Table 5.** Affiliated organisms found in the Mtl04, Mtl05 and Mtl06 sampling site by mtDNA sequencing.

| Mtl10 | % | Mtl11 | % | Mtl12 | % |
| --- | --- | --- | --- | --- | --- |
| *Homo sapiens* | 0.812 | *Homo sapiens* | 6.45 | *Homo sapiens* | 1.45 |
| *Rhomboplites aurorubens* | 0.0018 | *Bos taurus* | 3.25 | *Bos taurus* | 0.113 |
| *Ondatra zibethicus* | 0.081 | *Sus scrofa* | 3.21 | *Ondatra zibethicus* | 3.02 |
| *Procyon lotor* | 0.0035 | *Gallus gallus* | 2.36 | *Castor canadensis* | 0.779 |
| *Anas platyrhynchos* | 0.967 | *Felis catus* | 0.299 | *Sciurus carolinensis* | 0.132 |
| *Branta canadensis* | 0.0044 | *Sciurus carolinensis* | 5.96 | *Chelydra serpentina* | 0.0510 |
| *Actitis macularius* | 0.0036 | *Ondatra zibethicus* | 0.116 | *Culaea inconstans* | 78.18 |
| *Lithobates catesbeianus* | 0.061 | *Procyon lotor* | 0.0048 | *Umbra limi* | 9.73 |
| *Anaxyrus* spp. | 0.0028 | *Sylvilagus floridanus* | 0.0015 | *Ameiurus nebulosus* | 3.93 |
| *Culaea inconstans* | 33.01 | *Zonotrichia* spp. | 0.363 | *Cyprinella spiloptera* | 0.770 |
| *Pimephales promelas* | 21.03 | *Branta canadensis* | 0.339 | *Esox lucius* | 0.725 |
| *Ameiurus nebulosus* | 13.65 | *Corvus* spp. | 0.207 | *Notemigonus crysoleucas* | 0.154 |
| *Umbra limi* | 1.08 | *Vireo* spp. | 0.151 | *Lepomis gibbosus* | 0.0498 |
| *Lepomis gibbosus* | 0.160 | *Larus* spp. | 0.138 | *Ictalurus punctatus* | 0.0229 |
| *Catostomus* spp. | 0.012 | *Anas platyrhynchos* | 0.133 | *Pimephales promelas* | 0.0113 |
| *Notemigonus crysoleucas* | 0.0086 | *Anaxyrus* spp. | 0.760 | *Catostomus* spp. | 0.0053 |
| *Phoxinus* spp. | 0.0040 | *Lithobates sylvaticus* | 0.0218 | *Plumatella* | 0.872 |
| *Esox lucius* | 0.0026 | *Pimephales promelas* | 46.92 | *Ilyocryptus* spp. | 0.0086 |
| *Cypridopsis vidua* | 29.05 | *Carassius auratus* | 22.35 | *Eristalis* spp. | 0.0019 |
| *Bosmina freyi* | 0.050 | *Culaea inconstans* | 1.60 |  |  |
| *Plumatella rugosa* | 0.0074 | *Lopholatilus chamaeleonticeps* | 0.354 |  |  |
| *Pseudendoclonium akinetum* | 0.0072 | *Sander vitreus* | 0.202 |  |  |
| *Corallina chilensis* | 0.0011 | *Ameiurus nebulosus* | 0.0215 |  |  |
| *Halteria grandinella* | 0.0003 | *Lepomis gibbosus* | 0.0099 |  |  |
|  |  | *Catostomus* spp. | 0.0075 |  |  |
|  |  | *Cyprinus carpio* | 0.0039 |  |  |
|  |  | *Esox lucius* | 0.0038 |  |  |
|  |  | *Notemigonus crysoleucas* | 0.0024 |  |  |
|  |  | *Plumatella* | 4.70 |  |  |
|  |  | *Bosmina freyi* | 0.0410 |  |  |
|  |  | *Melanoplus* spp. | 0.0204 |  |  |
|  |  | *Cypridopsis vidua* | 0.0048 |  |  |

...: human, pet and livestock/meat. ...: Wild mammals. ...: birds. ...: amphibians, reptile. ...: fish. ...: others such as cretaceans, insects.

### Mtl11 : Stream *De Montigny* (Table 5)

The stream is the outlet of a pond located in dense urban and commercial areas. The sampling site was adjacent to a busy boulevard. We found a high proportion of sequences from the small fish fathead minnow (*Pimephales promelas*; 46.9%), and the invasive fish goldfish (22.4%). Human sequences were found at 6.5%. Sequences affiliated with bovine (3.3%), porcine (3.2%), and poultry (2.4%), at unexpected relative levels, were found. Nearby, there are various food and meat stores around the pond. Sequences affiliated with the squirrel (6.0%) were found at a similar relative level to human sequences.

### Mtl12 : Marsh *parc-nature de la Pointe-aux-Prairies* (Table 5)

This stream flows through a municipal park, in a mostly wooded area, far from the urban area, except for golf courses and a nearby school. High proportions of sequences affiliated with the small fish Brook stickleback (78.2%) and Central mudminnow (*Umbra limi*; 9.7%) were found. Unlike the River *à l’Orme* site, the diversity of mammals, fish, and birds in this sample was low. However, due to the high proportion of stickleback sequences, the sequencing depth was probably not sufficient to observe other species. Sequences affiliated with humans (1.5%) and bovine (0.11%) were found, as well as muskrat (3.0%) and beaver (*Castor canadensis*; 0.78%).

## Discussion

Human-related mtDNA sequences were found in all water bodies. Human activities can generate occasional releases of organic elements of animal origin into surface waters, for example, through fecal contamination, or through the discharge in wastewater of meat or fish waste, in addition to gray water from showers and laundry. These contaminations can come from faulty septic tanks, municipal wastewater discharged into water bodies without treatment, overflows, or faulty connection of wastewater pipes. In addition, multiple animal species can be detected in human feces or wastewater (Caldwell *et al.*, 2007, Caldwell & Levine, 2009, Kortbaoui *et al.*, 2009). For instance, we demonstrated in a previous study that bovine mtDNA sequences were 10 to 50 times lower than human mtDNA sequences in municipal wastewater, and even less in porcine or poultry (Ragot & Villemur, 2022). These sequences probably originated either from incomplete digestion of meat by the human digestive tract or from kitchen waste ending up in wastewater. Thus, obtaining sequences affiliated with bovine, poultry or porcine, and even moose (Mtl05), or certain fish (Atlantic salmon Mtl04, vermillion snapper Mtl09) in association with sequences affiliated with human, could be due to their consumption, and their release in human feces/wastewater.

However, when mtDNA sequences affiliated with bovine, porcine, and poultry are found at relative levels similar to those of human, as observed at three sites (Mtl06, Mtl08, and Mtl11), other sources must be considered. Since no livestock farms were present nearby on the Island of Montreal, one possibility is the discharge of meat by commercial food stores into the sewers with defective wastewater pipes. Another possibility is the discharge by these stores of food waste directly into water bodies. This could be the case in Bouchard Creek, where sequences from several marine fish species were found in significant proportions.

The mtDNA method was able to detect invasive fish such as pumpkinseed, brown bullhead, and northern pike in several water bodies. However, we did not find sequences associated with exotic animals. In a previous study, we detected sequences associated with guppy (*Poecilia reticulata*) in the L’Assomption River (Ragot & Villemur, 2022). Our results suggest that the presence of exotic animals in the monitored water bodies is not significant. However, as mentioned before, the location of the water collection site in some water bodies may not have been appropriate for detecting these animals, as was the case at site Mtl09.

High proportions of crustacean-related sequences were obtained, as observed in samples Mtl03 (*Bosmina*, 65%), Mtl07 (*Daphnia* crustaceans, 36%), and Mtl10 (*Cypridopsis vidua*, 29%). Although the primers were not designed for these organisms, their abundance appears sufficient for them to pair with their mitochondrial sequences. However, the pairing of the primers with these sequences during PCR was likely suboptimal, making the accuracy of their proportion in the samples questionable. Moreover, their identification also remains questionable because the sequences related to the lineages of these organisms were generally less than 95% identical to known species. These are likely new species, or known species whose mitochondrial sequence has not yet been sequenced.

It is important to understand that the quantitative aspect of the sequencing method reveals proportions, not absolute concentrations, due to the nature of sequencing. We discussed this aspect in Villemur (2025). However, the magnitude of the proportion values could provide a good estimate of the release of organic matter by animals.

Although other studies monitored water bodies with mtDNA primers, mostly specific to 12S rRNA mtDNA sequences ((Palumbi *et al.*, 2002, Rasmussen *et al.*, 2009, Riaz *et al.*, 2011, Andersen *et al.*, 2012, Giguet-Covex *et al.*, 2014, Miya *et al.*, 2015), to my knowledge, no reports were found relative of monitoring water bodies in dense urban areas for profiling mammals, birds, fish and others by mtDNA sequencing. I recommend caution in the determination of the presence of specific animals as these animals may not be present at all as a living organism, but rather sequences originate from waste disposal.

## Conclusions

Our results showed that the mtDNA approach allows for the establishment of global profiles of mammals, birds, fish, amphibians, and reptiles present in water bodies and their surroundings. Invasive fish were easily detected, probably due to their abundance. In contrast, sequences associated with exotic animals such as turtles or coyotes were not found, probably due to their low numbers, but also due to the sampling location. For example, by sampling a fast-flowing river like the river *à l’Orme* (Mtl04), with significant water mixing, we were able to detect a very low proportion of animals (such as a black bear). Therefore, choosing the outlet of a pond with higher water mixing would be preferable (if possible). In addition, our approach allows for the detection of meat and fish waste discharges into water bodies. Our approach can therefore help the municipality to define precise objectives for the mitigation of these discharges, by assessing defective sanitation connections, or by identifying intentional discharges of waste into water bodies.

## Appendix Correspondence between the Latin names and the common names of the organisms described in the report

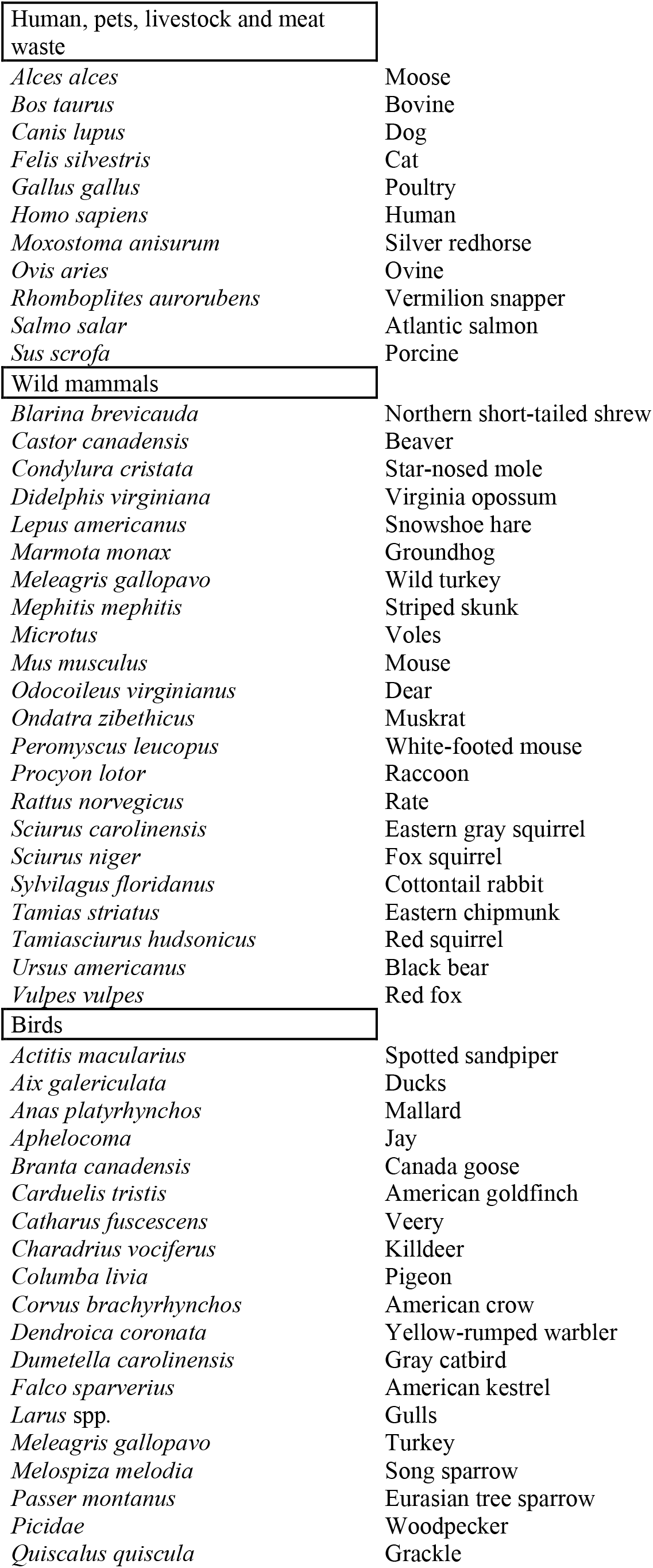
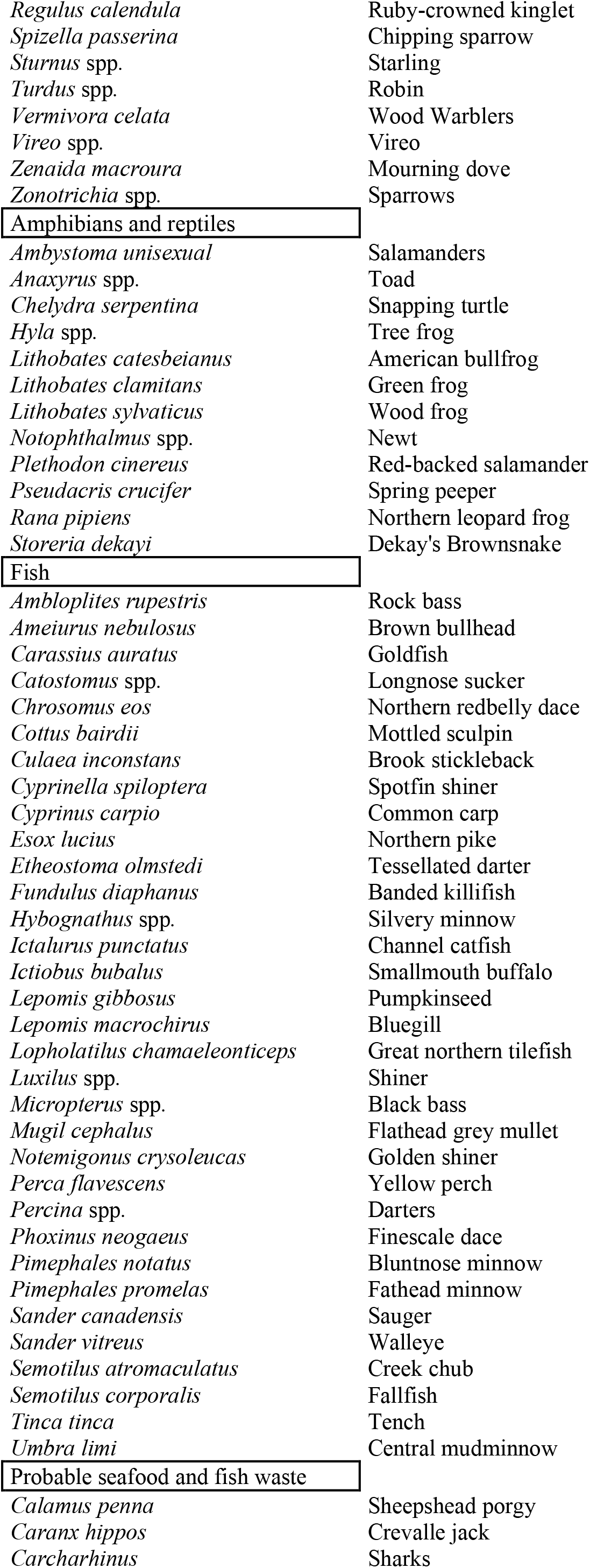
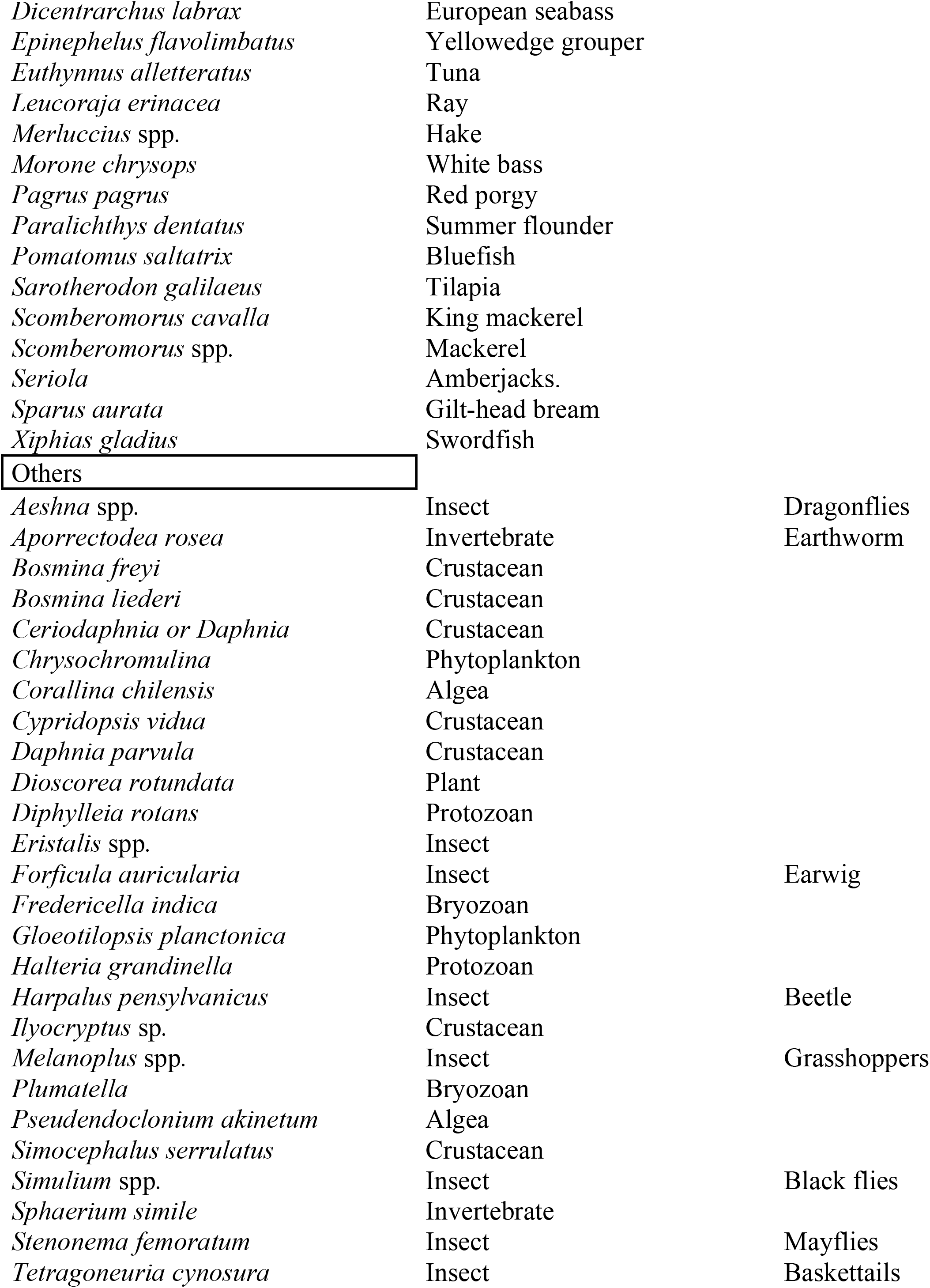

